# Exploring the coronavirus epidemic using the new WashU Virus Genome Browser

**DOI:** 10.1101/2020.02.07.939124

**Authors:** Jennifer A. Flynn, Deepak Purushotham, Mayank NK Choudhary, Xiaoyu Zhuo, Changxu Fan, Gavriel Matt, Daofeng Li, Ting Wang

## Abstract

Since its debut in mid-December, 2019, the novel coronavirus (2019-nCoV) has rapidly spread from its origin in Wuhan, China, to several countries across the globe, leading to a global health crisis. As of February 7, 2020, 44 strains of the virus have been sequenced and uploaded to NCBI’s GenBank [1], providing insight into the virus’s evolutionary history and pathogenesis. Here, we present the WashU Virus Genome Browser, a web-based portal for viewing virus genomic data. The browser is home to 16 complete 2019-nCoV genome sequences, together with hundreds of related viral sequences including severe acute respiratory syndrome coronavirus (SARS-CoV), Middle East respiratory syndrome coronavirus (MERS-CoV), and Ebola virus. In addition, the browser features unique customizability, supporting user-provided upload of novel viral sequences in various formats. Sequences can be viewed in both a track-based representation as well as a phylogenetic tree-based view, allowing the user to easily compare sequence features across multiple strains. The WashU Virus Genome Browser inherited many features and track types from the WashU Epigenome Browser, and additionally incorporated a new type of SNV track to address the specific needs of viral research. Our Virus Browser portal can be accessed at https://virusgateway.wustl.edu, and documentation is available at https://virusgateway.readthedocs.io/.

## Introduction

On December 12, 2019, the first case of a novel coronavirus (2019-nCoV) was reported in Wuhan, China, and by February 6, 2020, the virus spread to 24 additional countries, infecting more than 27,000 individuals and resulting in 565 fatalities, according to the World Health Organization (WHO) [2]. The 2019-nCoV is a member of the *Betacoronavirus* genus, which is one of four genera of coronaviruses of the subfamily Orthocoronavirinae in the family Coronaviridae, of the order Nidovirales [3, 4]. The species in this genus are enveloped, contain a positive single-stranded RNA genome, and are of zoonotic, likely bat, origins [5]. 2019-nCoV is one of the largest RNA virus genomes varying from 27kb to 32kb in size, with this particular strain ringing in at 29,903 bps long [6]. The virus is one of 7 coronaviruses known to infect humans, and along with the severe acute respiratory syndrome coronavirus (SARS-CoV) and the Middle East respiratory syndrome coronavirus (MERS-CoV), 2019-nCoV is one of the species responsible for severe respiratory distress in humans as well as other animals [4]. In an effort to better understand the pathogenesis of this family of viruses, several groups have sequenced individual strains, providing a powerful resource hosted by NCBI.

The WashU Epigenome Browser is a powerful tool for visualizing multiple functional genomic datasets and data types simultaneously [5–8]. The general layout of the Epigenome Browser displays the genome on the x-axis, and individual tracks encompassing many different varieties can be loaded and viewed in the context of the genome and accompanying metadata. Recent updates to the browser have incorporated new functionality, including live browsing, greatly enhancing its functionality [5]. With this powerful tool in-hand, we sought to adapt the browser for use of visualizing viral genomes, to support more efficient research and more rapid knowledge dissemination in response to the recent 2019-nCoV outbreak. To accomplish this, we created the WashU Virus Genome Browser, adapted from the WashU Epigenome Browser. The Virus Genome Browser houses reference genomes for 2019-nCoV, MERS, SARS, and Ebola virus, along with several annotation tracks including gene annotation, putative antibody-binding epitopes, CG density, and sequence diversity. Complete genomes of individual strains of each virus species (16, 551, 332, and 1574, respectively as of February 7, 2020, and periodically updated) are available as a database for instant viewing on the Virus Browser via multiple track types designed to display pairwise comparison to the references. Additionally, we aligned the genomes of all available strains in the database and generated a phylogenetic tree for each virus species that allows the user to directly select strains from the tree and view as tracks in the genomic display. In addition to all track types supported by the Epigenome Browser, we designed a new SNV track type to display sequence variation. Users can upload their own alignment results from any aligner and display them as SNV tracks on the browser.

The functionality of the Virus Browser is not limited to the 4 species currently housed. Users can upload their own reference genome in FASTA format and display tracks in the context of the user-specified reference. While maintaining the same functionality as that of the Epigenome Browser and providing novel functionality to aid specifically in viral genome research, we hope that the Virus Browser may facilitate research against new epidemic viruses.

## Materials and Methods

### Reference sequences, additional strains, and gene annotations

Genomic sequences of all viral strains were downloaded as FASTA files from NCBI [Supplementary Table 1]. All available sequences as of January 31, 2020, for 2019-nCoV, MERS, SARS, and Ebola were downloaded (n=16, 551, 332, and 1574, respectively). The reference genomic sequence of the selected virus (2019-nCoV: NC_045512.2; MERS: NC_019843.3; SARS: NC_004718.3, Ebola: KM034562.1) is automatically displayed as a color coded track when opening the genomic track browser viewing format. Genic annotations of reference genomes were downloaded as GFF3 files from NCBI and converted to refBed format for viewing on the browser.

### Sequence alignment and tree generation

The genomes of all individual strains of each virus were aligned to the reference genome using the pairwise alignment tool stretcher [9] with parameters “-gapopen 16 -gapextend 4”. To generate the phylogenetic trees, we used the MAFFT program, employing the fast option to align individual strains of each viral genome to its reference [10, 11]. Phylogenetic trees were built using FastTree with the GTR model [12, 13].

### Data Tracks

#### Genome Comparison Track

We adopted the genome comparison tracks from the WashU Epigenome Browser. Any pairwise alignment results in markx3 or FASTA format can be converted with our publicly accessible script “aligned_fa_2_genomealign.py” [14] and directly displayed as genome comparison tracks on the Virus Browser.

#### SNV Track

We developed the SNV track type to display sequence variation of individual strains relative to their reference. Variations from the reference genome, including mismatches and deletions, are displayed with customizable colors. Insertions compared to the reference genome can be expanded upon selecting to show the nucleotides inserted. When viewing large regions, such as the whole genome, it is not possible to display all individual variation events. Therefore, the frequency of variation events is also displayed in a “density mode” where a high value over a region signifies multiple sequence variation events within the region.

#### Congeneric (or Closely-related) Immune Epitope Locations

We wrote a text processing utility to import antibody-binding epitopes curated by the Immune Epitope Database and Analysis Resource (IEDB) for MERS-CoV and SARS-CoV [15]. Subsequently, we used tblastn to align linear epitopes to the Wuhan seafood market pneumonia virus isolate Wuhan-Hu-1 (Taxonomy ID: 2697049; NCBI:txid2697049). We found 955 out of 2,817 linear epitopes identified in SARS had at least 1 “hit” in the 2019-nCoV genome [Supplementary Data 1]. Three epitopes have 2 “hits” each. However, the secondary hit is on the negative strand with very low percent identity (37.5% to 53.8%) to the 2019-nCoV genome and are hence filtered out as 2019-nCoV is a (+) ssRNA virus. Similarly, we found 1 hit out of 38 linear epitopes identified in MERS. We also provide scripts [14] that can be used to obtain a quick overview of the similarity of linear epitopes identified in other viruses in databases like IEDB. These tracks can provide researchers preliminary data to support exploratory analyses pertaining to the immunogenicity of 2019-nCoV—an actively explored vertical of 2019-nCoV research.

#### GC Density Track

GC density tracks were created for each reference genome, displaying the percentage of G (guanine) and C (cytosine) bases in 5-bp windows.

#### Sequence Diversity Track and Shannon Track

In order to display a measure of sequence conservation across the genome, we calculated the percentage of each of the 4 nucleotides at each position in the genome across all strains for a given virus species. The resulting bed tracks display the percentages each nucleotide comprises across all strain for each genomic position. We also calculated Shannon entropy for each position along the genome using the percentages of each of the 4 nucleotides. A high Shannon entropy at a position signifies that the 4 possible nucleotides are equally likely across all strains of this virus, and thus the position is likely divergent. A low Shannon entropy at a position means that the identity of the nucleotide at this position is highly conserved across all strains. The entropy() function of the R package “entropy” was used for calculations.

#### Resources for User-Defined Bed and Categorical Tracks

In addition to our housed data tracks, we also offer scripts (“publicParseAlignment.py”, “publicAlignment.py”, and “publicConvertMarkx3.py”) to convert any markx3 or FASTA-formatted alignment into displayable bed and categorical formats, and a script (“publicJsonGen.py”) to generate a json file for uploading multiple data files together for display [https://github.com/debugpoint136/WashU-Virus-Genome-Browser]. A default color code for sequence variation is also included in the script.

## Results

### Organization of the Virus Genome Browser

The WashU Virus Genome Browser houses consensus reference genomic sequences for 4 different pathogenic virus species: 2019-nCoV, MERS, SARS, and Ebola, as well as a comprehensive set of genome assemblies for the individual strains of each virus (16, 551, 332, and 1574, respectively). When users first navigate to the WashU Virus Browser and select “Browse Data”, they are directed to a page with several customizable options, including a drop-down menu from which they may choose a reference genome [Figure 1]. Corresponding with the reference genome selected, a metadata table is displayed containing sortable features such as species, strain, isolate, isolation source, host, country, and collection date, to allow for quick and easy sorting of individual strains. The user may select viral isolates from the metadata table to be visualized in one of our two displayable platforms: the track view (green arrow, Figures 2 and 3) or the phylogenetic tree view (orange arrow, Figures 4 and 5).

**Figure 1:**
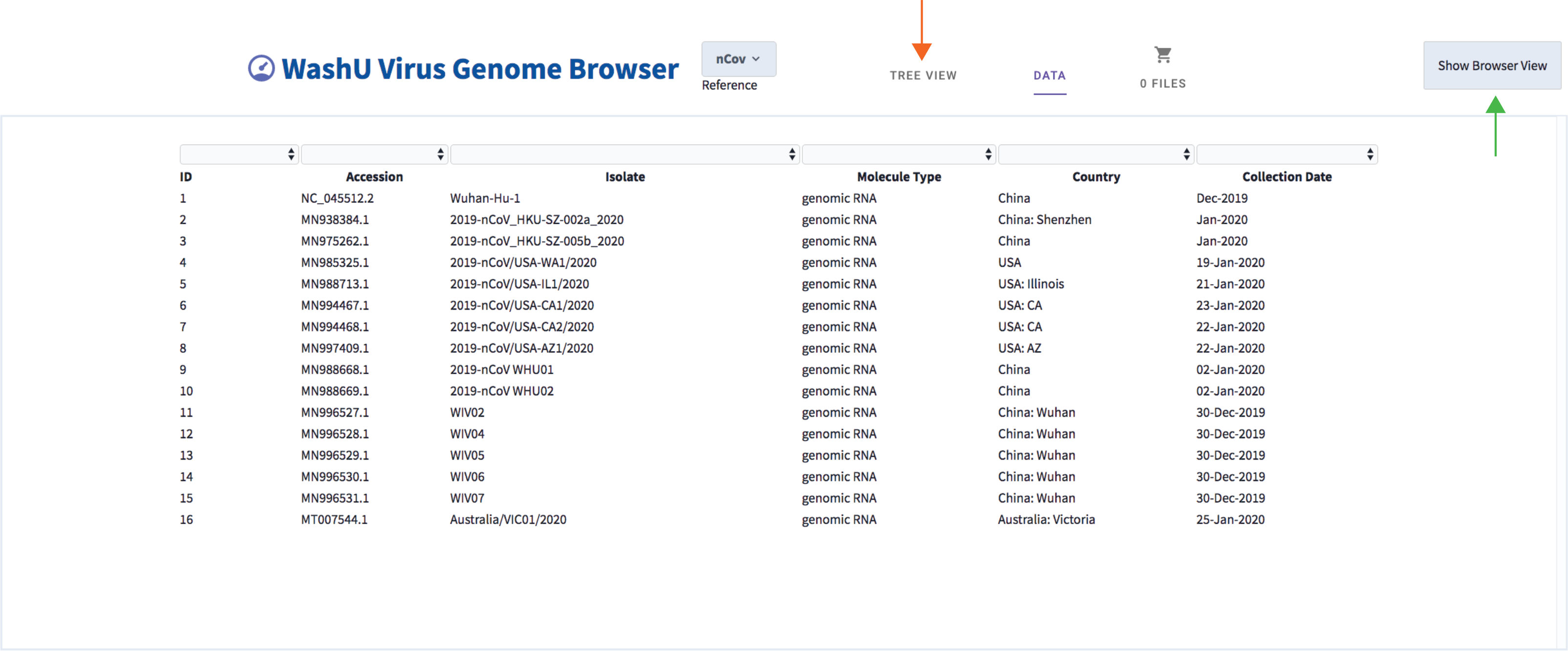
Screenshot of the WashU Virus Genome Browser data page. This view demonstrates several customizable features of the browser, including which genome reference to use, which data tracks to select based on several metadata features, and which browser view to use: “genomic” view (green arrow) or phylogenetic tree view (orange arrow).

### The Track View

The track view option has a standard genome browser layout similar to that of the WashU Epigenome Browser, in which a reference genome sequence is visualized as a sliding window. Various annotation data tracks are hosted on the browser and can be loaded for visualization in a genomic context. For each virus, we downloaded publicly available annotations of the reference genome and converted these annotations into refBed tracks that can be visualized in the genome browser. Likewise, immune epitopes identified in SARS were aligned to the 2019-nCoV reference [Materials and Methods], and a track displaying their coordinates in 2019-nCoV is provided. GC-density tracks were also created for each reference genome, and display the percentage of Gs (Guanines) and Cs (Cytosines) per 5bp window. An entropy track [Materials and Methods] showing the degree of sequence diversity at each position and a diversity track [Materials and Methods] showing the percentage of each of the 4 nucleotides at each position across all strains of the given virus species are also included in the database. In addition to hosting 4 virus species reference genomes, The Virus Genome Browser also supports displaying user-specified genomes provided in FASTA format, as shown in the top left part of Figure 2A, under the browser logo.

**Figure 2:**
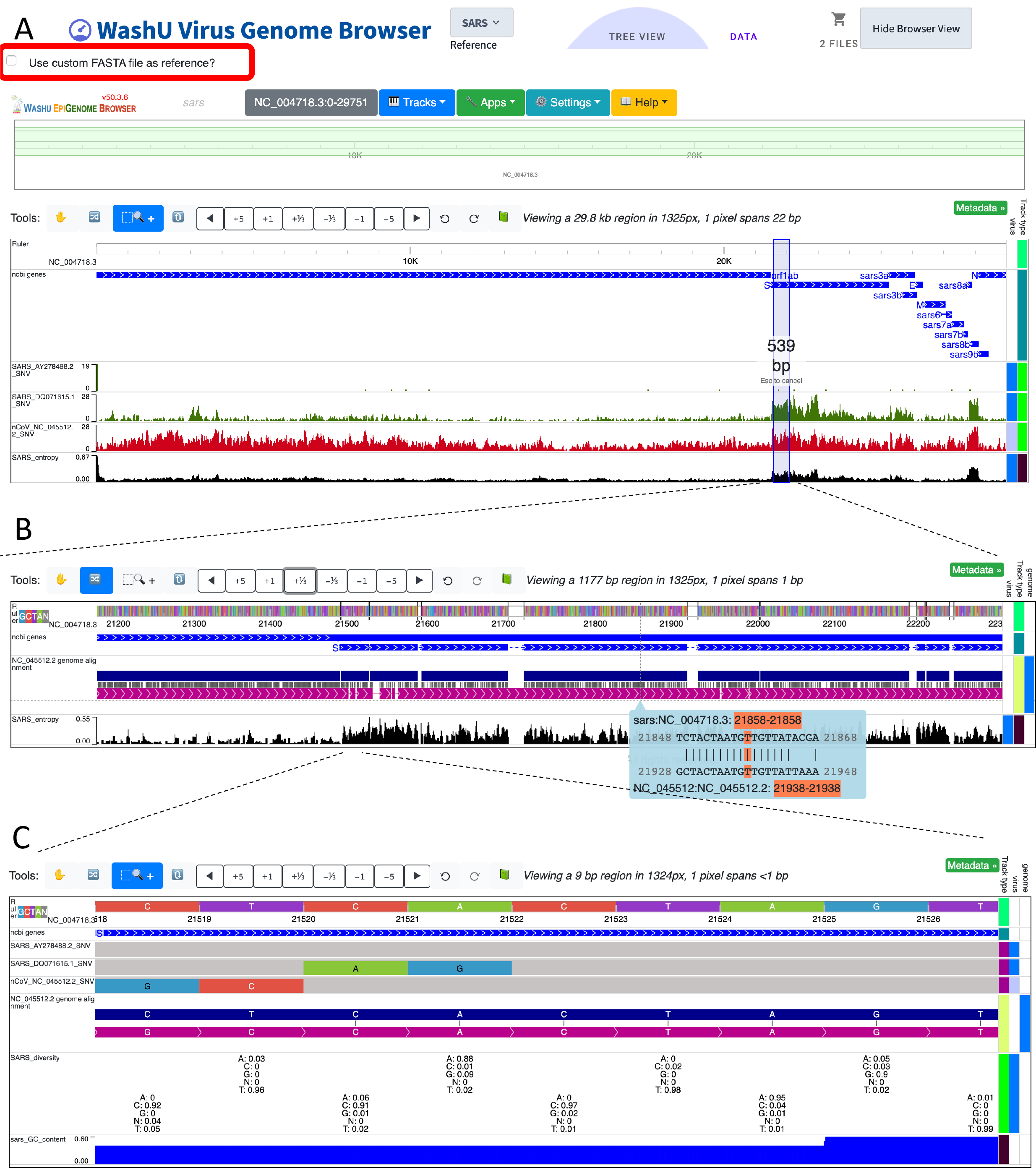
Illustration of genomic-level and nucleotide-level track views. A: “zoomed out” track view of the entire genome. 2019-nCoV reference genome (shown in red, NC045512.2) and 2 SARS strains (shown in green, DQ071615.1 and AY278488.2) are aligned to the SARS reference genome (NC_004718.3). The box in the top left corner allows users to upload and use any sequence in FASTA format as the reference genome. The shaded vertical bar demonstrates the user’s ability to select a region by mouse for further magnification. B: “Zoomed in” view of the sequence flanking the 5’ end of the S protein. C: A further “zoomed in” view to the level of individual nucleotides. Stretches of grey indicate matching while variations are color coded.

The WashU Virus Browser supports a “zoomed-out” view of the entire viral genome. The zoomed-out view can help the user quickly determine the regions of interest that have high frequencies of variation from the reference (SNV track), and also the regions with high nucleotide diversity among all strains (Shannon tracks) [Figure 2A]. Figure 2A illustrates a genome-level browser view of the 2019-nCoV reference genome and 2 SARS strains, each aligned to the SARS reference genome (AY278488.2 = BJ01, DQ071615.1 = Bat rp3, NC_045512.2 = 2019-nCoV). Sequence variation displayed in density mode [Materials and Methods] shows that the divergence between the 2019-nCoV reference genome (red) and the SARS reference genome is higher than the divergence between the two additional SARS strains (green) and the SARS reference genome. For AY278488.2, the variation from reference is mainly confined to the beginning of the genome, while the remainder of the genome is relatively consistent with the reference. However, for DQ071615.1 (bat-derived), the 5’ end of gene S displays high variation from the reference genome. Likewise, the SARS Shannon track shows that the SARS genome is highly diverse across different strains at gene S.

Once a region of interest is identified, the standard magnification tool of the browser can be used to quickly zoom into the region [Figure 2A]. Upon zooming in, a genome comparison track can be used to inspect variations from the reference genome, particularly useful for comparing cross-species alignments and viewing structural variations [Figure 2B]. The genome comparison track is adopted from the Epigenome Browser. The top navy-colored horizontal bar represents the reference genome loaded (SARS in the case of Figure 2B) and the bottom purple-colored horizontal bar represents the sequence being aligned to the reference (the 2019-nCoV reference sequence, NC_045512.2, in this case). Insertions and deletions are represented as gaps in either the reference or the query. Matches are represented by black lines linking the 2 genomes while mismatches are distinguished by omission of the black bar. When the user hovers over a specific nucleotide, the alignment details around that specific nucleotide are shown.

Upon further magnification, regions can be inspected on a nucleotide level. Mismatches, insertions, and deletions are color-coded in the SNV tracks and stretches of grey signify positions matching the reference [Figure 2C]. Detailed information, such as inserted nucleotides, is displayed upon clicking. When zoomed into individual nucleotides, as shown in Figure 2C, The diversity bed track shows the percentage of each nucleotide across all strains of SARS at the specific position.

The versatility of the WashU browser framework makes it possible to adapt the browser to address various questions of interest. Figure 3 demonstrates the utility of using the browser for immune epitope conservation discovery. We recapitulated Zhou et al.’s [16] alignment results of two SARS strains to the reference 2019-nCoV nucleocapsid protein sequence [Figure 3A, 3B]. Upon inspection of the region, we could directly observe that many immune epitopes are conserved between SARS and 2019-nCoV [Figure 3C]. The user can identify the amino acid sequence of an epitope by simply clicking the track.

**Figure 3:**
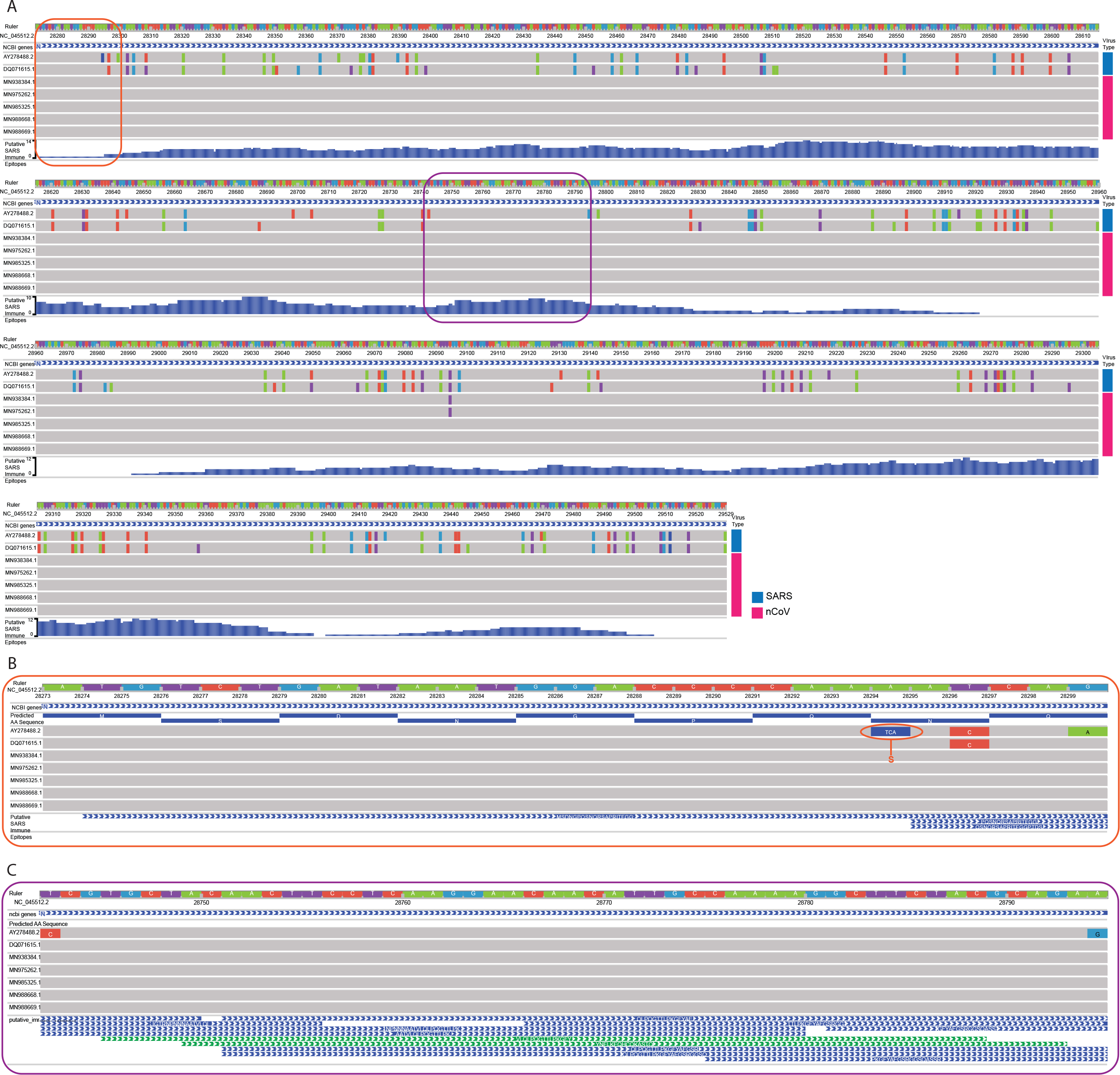
Alignment of the genomic region encoding the nucleocapsid protein. A: 2 SARS strains (DQ071615.1 and AY278488.2) and 5 2019-nCoV strains (MN938384.1, MN975262.1, MN985325.1, MN988668.1, and MN988669.1) are aligned to the 2019-nCoV reference. The region encoding the nucleocapsid protein is shown. Putative SARS immune epitopes [Materials and Methods] are displayed in “density mode”. B: A zoomed-in view of A (orange box), displaying the first 9 amino acids of the reference. Results show a “TCA” insertion in the AY278488.2 alignment between positions 28294 and 28295 of the 2019-nCoV reference sequence, which is not present in DQ071615.1. These results are consistent with the results reported in Extended Data Figure 5 of Zhou et al. [16]. C: A zoomed-in view of A (purple box), displaying a region conserved between SARS and 2019-nCoV, overlapping several putative immune epitopes.

Encouraged by the high sequence similarity between SARS-CoV and the 2019-nCoV reference strain (NCBI:txid2697049), we mined the list of experimentally identified linear epitopes from T-cell, B-cell and MHC-ligand assays from IEDB [15]. We identified a list of 320 high-confidence linear epitopes [Supplementary Table 2] whose amino acids are identical to predicted translated products from the 2019-nCoV reference strain. These provide a catalogue of epitopes for researchers testing immune targets that can potentially elicit T-cell, B-cell and antibody response to 2019-nCoV.

We also provide these as an annotated bed track to the reference 2019-nCoV genome. Along with the individual strains’ SNV tracks, the epitope tracks can provide a quick, intuitive and visual resource to guide prioritization of experimental resources towards developing diagnostics and therapeutics against 2019-nCoV. The value of our novel SNV tracks will only increase as additional strains are sequenced, helping us better understand the evolving 2019-nCoV genome and prioritize epitopes.

### The Phylogenetic Tree View

The second viewing option offered by the WashU Virus Genome Browser is a “tree” format, in which the evolutionary relationships of different viral isolates can be visualized as a phylogenetic tree [17]. When the user navigates to the data page of the browser, and selects “Tree View” [Figure 1], all viral genomes hosted on the browser for the selected virus species are displayed in the form of a right-aligned phylogenetic tree, where solid lines indicate branch lengths [Figure 4]. To the right of the tree is a metadata heatmap displaying strain-specific details such as isolate, isolation source, host, country, and collection date. Additionally, if the user added any individual tracks to their cart from the main page, those selected will display a checkmark to the right, allowing the user to easily see where their strains of interest lie among all other strains.

**Figure 4:**
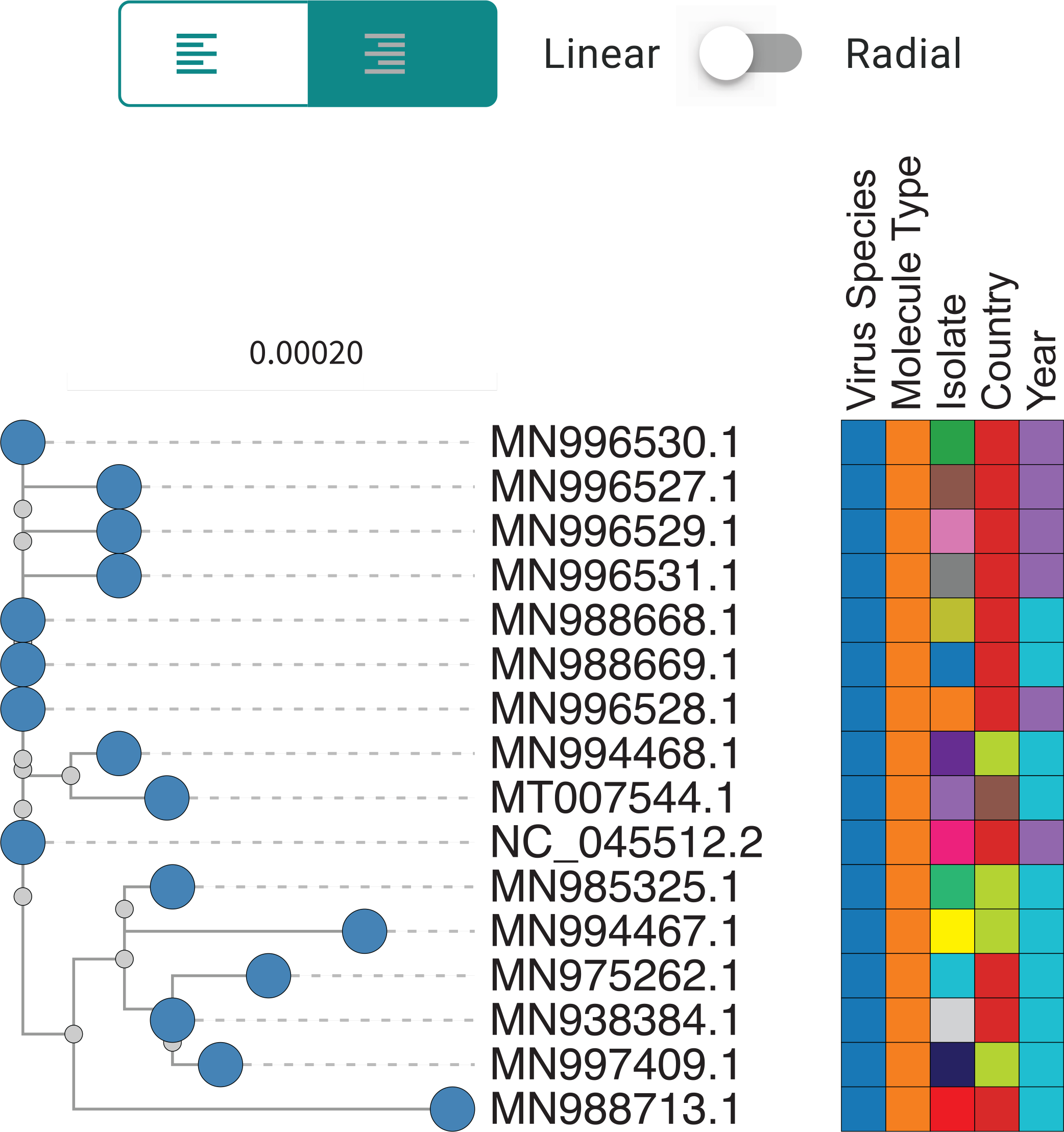
Screenshot of a linear, right-aligned tree view displaying all housed 2019-nCoV sequences with accompanying metadata. Solid lines signify distance.

In addition to the right-aligned tree view, the browser also supports a more traditional left-aligned linear tree view and a radial view. The left-aligned tree view displays branch lengths indicating relatedness of isolates [Figure 5A]. We noticed that in each virus type, several individual strains maintained high sequence similarity, resulting in several short branch lengths and a long vertical tree. In order to improve visualization, we also created a radial tree view [Figure 5B].

**Figure 5:**
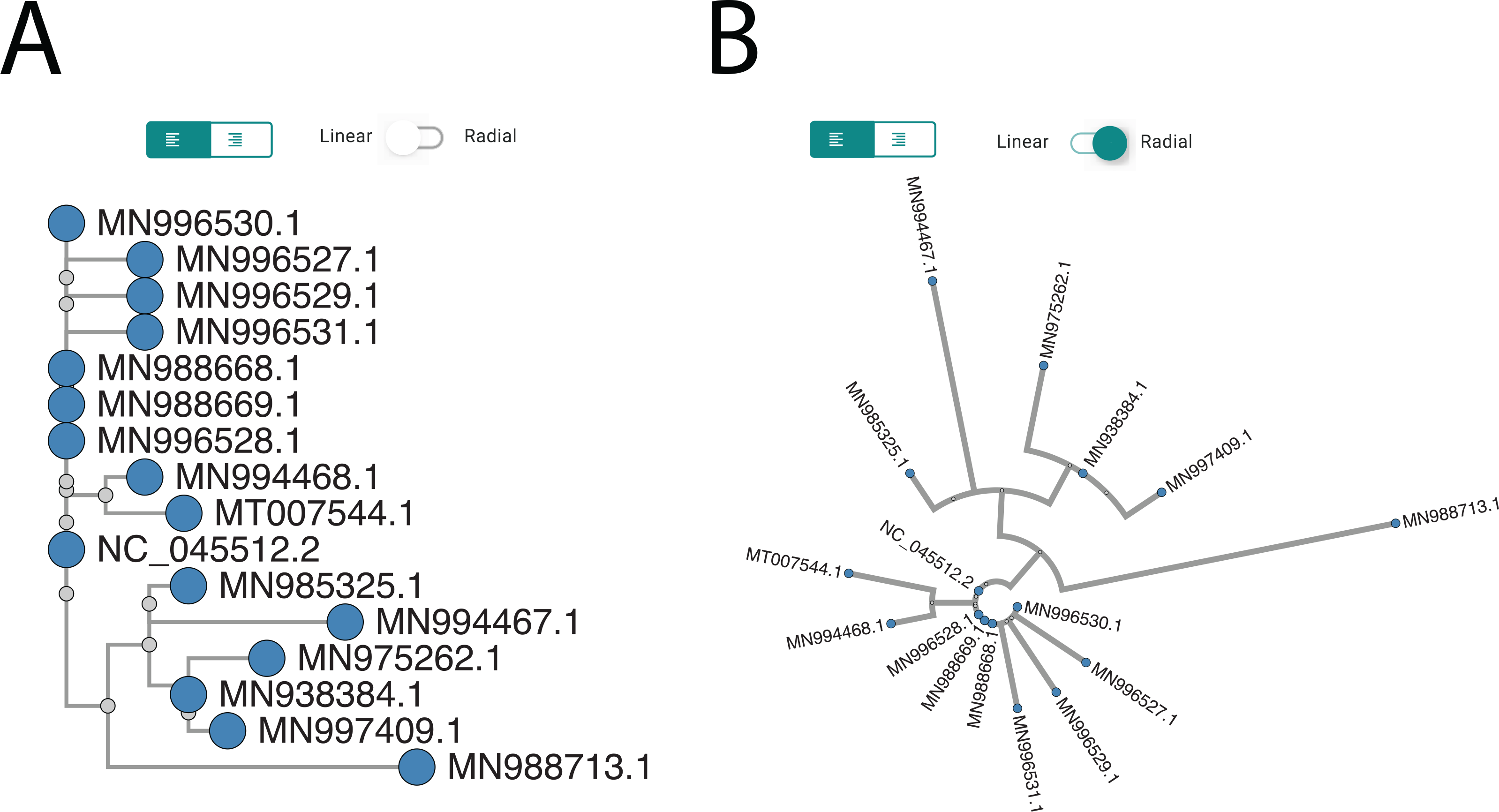
A: Screenshot of a linear, left-aligned phylogenetic tree view, displaying all 2019-nCoV strains hosted by the browser. B: Screenshot of a radial tree view for all 2019-nCoV strains.

## Discussion

Maps help us understand the world around us and navigate it. Moreover, they play a critical role in disaster management during disease outbreaks. Herein, we describe the first genetic mapping, exploration, and visualization tool from the WashU Epigenome Browser team that is specifically dedicated to viral genomes. We provide reference genome maps and genomic datasets related to 4 viral disease outbreaks: SARS (2002-03), MERS (2012), Ebola (2014-16) and the latest nCoV (2019-20). More importantly, we not only present publicly available information in the format of easily accessible data tracks, but also offer a platform with high customizability and flexibility where individual investigators and teams can upload and visualize their own genomic datasets in a plethora of formats. In this report, we have demonstrated using the Virus Browser to 1) quickly and intuitively compare multiple viral genomes and study the viral genome at multiple levels [Figure 2, Figure 4, Figure 5]; and 2) combine viral genome information with other functional genomic information (amino acid sequence and putative immune epitope locations, as shown Figure 3) through multiple track types the browser supports, and identify potential therapeutic targets.

We expect that the WashU Virus Browser can support research related to the latest novel Coronavirus outbreak of 2019-20, and hope that this tool helps accelerate research to further our understanding of 2019-nCoV and aid in the development of therapeutics. In addition, our platform supports the study of any user-specified viral genome, and can be expanded to other viral research.

To aid in the battle against this crisis, we are releasing the browser at first moment. The browser is still under active construction and is constantly being updated. General feedback, suggestions for additional tracks, and bug reports may be sent to the WashU Virus Genome Browser team by opening an issue request at https://github.com/debugpoint136/WashU-Virus-Genome-Browser/issues.

## Acknowledgements

We thank doctors, nurses, investigators, and all other people fighting on the front line against this viral outbreak, and we sincerely hope that this tool will aid in this battle.

## Author Contribution

Conceptualization, T.W. Web development, D.L and D.P. SNV track development, J.F. and C.F. Immune epitope analysis, M.C. Data download, metadata generation and annotation, G.M. Sequence alignments and tree generation, X.Z. Manuscript preparation, J.F, C.F, M.C, G.M, T.W.

